# Time-resolved FRET screening identifies small molecular modifiers of mutant Huntingtin conformational inflexibility in patient-derived cells

**DOI:** 10.1101/2021.01.18.427095

**Authors:** Johannes H. Wilbertz, Julia Frappier, Sandra Muller, Sabine Gratzer, Walter Englaro, Lisa M. Stanek, Barbara Calamini

**Author notes:** Ksilink, 16 rue d’Ankara, 67000 Strasbourg, France (Johannes H. Wilbertz).

## Abstract

Huntington’s disease (HD) is the most common monogenic neurodegenerative disease and is fatal. CAG repeat expansions in mutant Huntingtin (mHTT) exon 1 encode for polyglutamine (polyQ) stretches and influence age of onset and disease severity, depending on their length. mHTT is more structured compared to wild-type (wt) HTT, resulting in a decreased N-terminal conformational flexibility. mHTT inflexibility may contribute to both gain of function toxicity, due to increased mHTT aggregation propensity, but also to loss of function phenotypes, due to decreased interactions with binding partners. High-throughput-screening techniques to identify mHTT flexibility states and potential flexibility modifying small molecules are currently lacking. Here, we propose a novel approach for identifying small molecules that restore mHTT’s conformational flexibility in human patient fibroblasts. We applied an antibody-based time-resolved Förster resonance energy transfer (TR-FRET) immunoassay, measuring endogenous HTT flexibility using two validated HTT-specific antibodies. The ratio of TR-FRET signal at 4°C and 20°C differs between wtHTT and mHTT and allowed to perform a high-throughput screening using HTT flexibility as a read-out. We identified several small molecules that can partially rescue mHTT inflexibility, presumably by altering HTT post-translational modifications. This novel screening approach has the potential to identify previously unknown HD drugs and drug targets.

## Introduction

Huntington’s disease (HD) is the most common monogenic neurodegenerative disease affecting between 3-14 individuals per 100,000 (1,2). HD manifests itself by involuntary movements, cognitive deficits, impaired speech and swallowing, and eventually leads to death. Ages of onset and disease severity are inversely correlated with the length of the CAG repeat expansion encoding polyglutamine (polyQ) stretches within exon 1 of the Huntingtin *(HTT)* gene. Although the precise molecular mechanism for mutant HTT (mHTT) toxicity is unknown, there is substantial evidence that the polyQ expansion in mHTT causes a toxic gain-of-function phenotype (3). However, several gene dosage-modifying studies have also shown that the polyQ expansion within mHTT can also cause loss-of-function effects. wtHTT has been shown to interact with a large number of effector proteins. As a consequence, loss of normal HTT function can alter the activity and cellular localization of different partners thus having a negative impact on cellular physiology.

Despite advances to lower the cellular concentration of mHTT by antisense oligonucleotides, certain technical challenges and medical risks remain (4,5). wtHTT’s physiological function is still not fully understood and reducing both the mutant and wildtype (wt) HTT alleles might come at the risk of neurodevelopmental defects (4,6–10). Asymptomatic mHTT gene carriers, who in principle could benefit the most from reduction in mHTT levels, might also be at a higher risk for detrimental neurological sideeffects. Furthermore, the intrathecal administration of ASOs is a burdensome procedure for HD patients. ASOs are a relatively new type of treatment and it is not yet clear what collateral effects they may cause. Therefore, alternative modalities that complement or eventually replace mHTT lowering approaches might become fruitful strategies to treat HD.

A long standing question in HD research has been why a polyQ tract beyond 37 repeats results in pathogenesis. Recent studies suggest that the normal polyQ tract in wtHTT is mostly unstructured and functions as a critical conformational hinge that mediates interactions between wtHTT flanking protein-protein interaction domains (11,12). However, when the polyQ domain reaches the 37Q threshold, a reduced flexibility of the hinge region is observed (13–16). As the conformational dynamism is likely to be essential in modulating a range of intra- and inter-molecular interactions, when abnormal, it can lead to altered HTT function and disease pathogenesis. Multiple studies have shown that post-translational modifications (PTMs) in the HTT N17 domain, in particular phosphorylation at threonine 3 (T3) and at serines 13 and 16 (Ser13 and Ser16), can restore mHTT conformational flexibility. This effect is associated with reduced mHTT aggregation and cellular toxicity (14,17–20). These findings are further supported by HD mouse models in which N17 phosphomimetic mutations have been introduced and which demonstrate an *in vivo* prevention or reversal of mHTT-induced neuronal toxicity (17,21). Importantly, these findings indicate that HTT protein conformation is amenable to small molecule-induced modulation since PTMs are controlled by enzymes.

The enzymes responsible for N17 phosphorylation are unknown, but a few molecules that possess N17-modifying capabilities have been identified and were shown to rescue aggregation behavior and toxicity (21–23). However, the currently known small molecule modifiers of HTT PTMs have not been clearly linked to a mHTT flexibility rescue. In addition, these compounds do not all display drug-like properties.

Therefore, in order to identify novel modulators of mHTT PTMs, we developed a high-throughput screening approach in HD patient fibroblasts expressing endogenous levels of HTT. We utilize a previously described time-resolved Förster Resonance Energy Transfer (TR-FRET) immunoassay to detect and quantify HTT conformational flexibility changes (14–16). The presented screening approach is sensitive enough to measure HTT conformational flexibility changes in human fibroblast cells expressing endogenous mHTT with polyQ expansions present in the majority of the HD patient population (< 60 Qs) (24–28) and is amendable to high-throughput testing. As the PTM status of mHTT can play a central role in HD pathogenesis by influencing mHTT conformational flexibility, aggregation and toxicity, we hypothesize that compounds that modulate mHTT PTMs could provide a promising avenue for novel HD therapeutics.

## Results

Here, we use a previously described, scalable, and sensitive TR-FRET-based immunoassay assay, which can faithfully detect conformational flexibility differences between wtHTT and mHTT in response to variations in temperature (14–16). HTT conformational changes can be assessed by the relative positions of two antibodies labeled with acceptor and donor fluorophores, which recognize specific epitopes of the target protein: the N-terminal 2B7 antibody, which recognizes both wtHTT and mHTT, and the mHTT (polyQ-specific) MW1 antibody (Figure 1A). This assay provides a useful screening tool to identify small molecule modulators of mHTT conformational flexibility.

**Fig 1:**
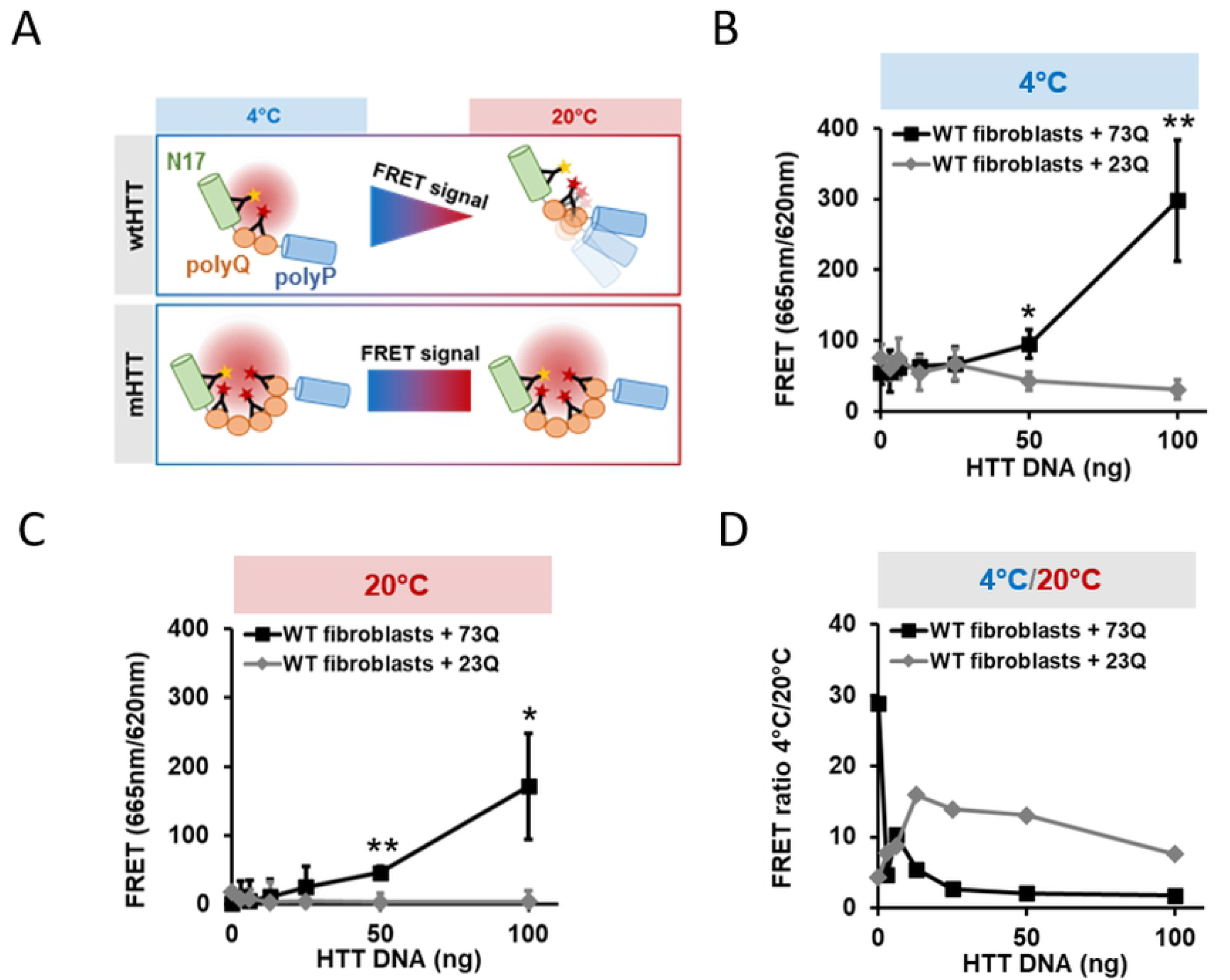
TR-FRET can distinguish transfected wtHTT and mHTT conformational flexibility in complex cell lysates. (A) wtHTT TR-FRET signals decrease upon temperature shifting from 4°C and 20°C, while mHTT signals stay relatively constant due to differing protein flexibility and epitope accessibility. (B) and (C) 73Q HTT or 23Q HTT transfected into wtHTT expressing human fibroblasts can be distinguished from each other both at 4°C and 20°C in complex cell lysates in a dose-responsive manner. Statistical comparison between transfection conditions at equal DNA concentration. (D) The 4°C/20°C TR-FRET ratio is higher for 23Q HTT than 73Q due to the stronger signal decrease at 20°C. All data are represented as mean ± S.D. of at least three independent experiments. Statistical significance was calculated using the paired two-tailed Student’s t-test (*=p<0.05; **=p<0.01; ***=p<0.005).

### TR-FRET can distinguish HTT flexibility states

The presented assay is able to detect conformational constraints imposed by polyQ expansion on HTT in lysates obtained from immortalized human HD fibroblasts, where HTT is expressed at endogenous levels from the relevant genomic locus (14). Screening for potentially PTM modifying small molecules must be performed in living cells in order to guarantee the integrity of all signaling pathways that can alter PTMs. Importantly, within cells antibody/epitope binding can be disturbed by non-specific interactions leading to unexpected or lowered FRET signals. In order to test whether we could accurately distinguish between the conformational flexibility of wtHTT and mHTT using the TR-FRET immunoassay in cell lysates, we transfected DNA plasmids containing wtHTT (23Q) or mHTT (73Q) DNA into human healthy donor-derived fibroblasts. This experimental setup guaranteed an isogenic background for testing the effect of temperature on the flexibility of overexpressed wtHTT and mHTT proteins. After 48 hours of construct expression, we performed the TR-FRET assay on the cell lysates and measured the signal generated by proximity of the 2B7-Tb and MW1-D2 antibody pair at both 4°C and 20°C HTT. We hypothesized that the 4°C/20°C TR-FRET signal ratio generated by the wtHTT protein would be higher than that of mHTT due to wtHTT’s increased flexibility at 20°C (Fig 1 A). In addition, we expected that the absolute TR-FRET signal should be higher for mHTT at both 4°C and 20°C due to the presence of higher number of polyQ epitopes, which increase the avidity of the MW1-D2 antibody for the mutant protein. As predicted, we observed that for increasing plasmid concentrations, the TR-FRET signal generated by the expression of mHTT was consistently higher than that generated by wtHTT (Fig 1 B and C). In addition, we also observed a decrease in total TR-FRET signal intensity for both constructs when cell lysates where shifted to 20°C. However, while the TR-FRET signal reduction for wtHTT was especially pronounced at this temperature and approached the detection limit (Fig 1 C), the signal for mHTT was just slightly lower that that obtained at 4°C. As a result, the 4°C/20°C TR-FRET ratio for wtHTT was consistently higher even at low plasmid concentrations than the temperature ratio of mHTT (Fig 1 D). Together these results indicate that the TR-FRET-based immunoassay is sensitive enough to detect specific HTT-type dependent conformational changes in complex cell lysates overexpressing wt and mHTT proteins.

Next, we tested whether our TR-FRET assay could detect endogenously expressed HTT proteins and whether the dual-temperature read-out was sensitive enough to detect flexibility differences between wtHTT and mHTT in cell lysates. We tested different primary HD patient fibroblast cell lines with varying polyQ lengths (S1 Fig A). The GM04691 fibroblast line displayed the highest signal-to-background (S/B) ratio and was selected for immortalization using hTERT + E6/E7 transduction (S1 Fig A). Most studies have performed TR-FRET assays on overexpressed HTT gene fragments containing very long (> 70) CAG repeats, which only occur in a minority of HD patients (24–26). The GM04691 cell line expresses endogenous 54Q mHTT and allowed us to investigate the properties of mHTT with a polyQ length much closer to the 40-50 Q repeat length most frequently observed in the HD patient population (27,28). Western blotting using a Tris-Acetate gel allowed the separation of both mHTT and wtHTT alleles expressed by GM04691 cells (S1 Fig B).

### mHTT flexibility can be modified in patient-derived human fibroblasts

In order to determine the optimal cell density to be used for obtaining the highest TR-FRET signal before reaching saturation, we performed the assay using different cell densities per well. We observed a clear cell concentration-dependent TR-FRET signal at 4°C for both wtHTT and mHTT (Fig 2 A). As expected, the absolute wtHTT signal was lower than the mHTT signal due to the lower number of polyQ epitopes for the MW1-D2 antibody present on the WT protein. At 20°C the fluorescence signals for both wtHTT and mHTT were reduced compared to the signal at 4°C (Fig 2 B). This effect was most likely related to the increased protein flexibility at higher temperatures yielding a lower FRET efficiency. When calculating the 4°C/20°C ratio, wtHTT showed a higher degree of flexibility than mHTT over all cell concentration ranges. An up to seven-fold difference was observed with a plateau effect reached at about 20-40×10^3^ cells/well, indicating that a high protein concentration and, consequently, a high number of cells per well is needed to obtain a robust signal ratio (Fig 2 C) with endogenously-expressed HTT.

**Fig 2:**
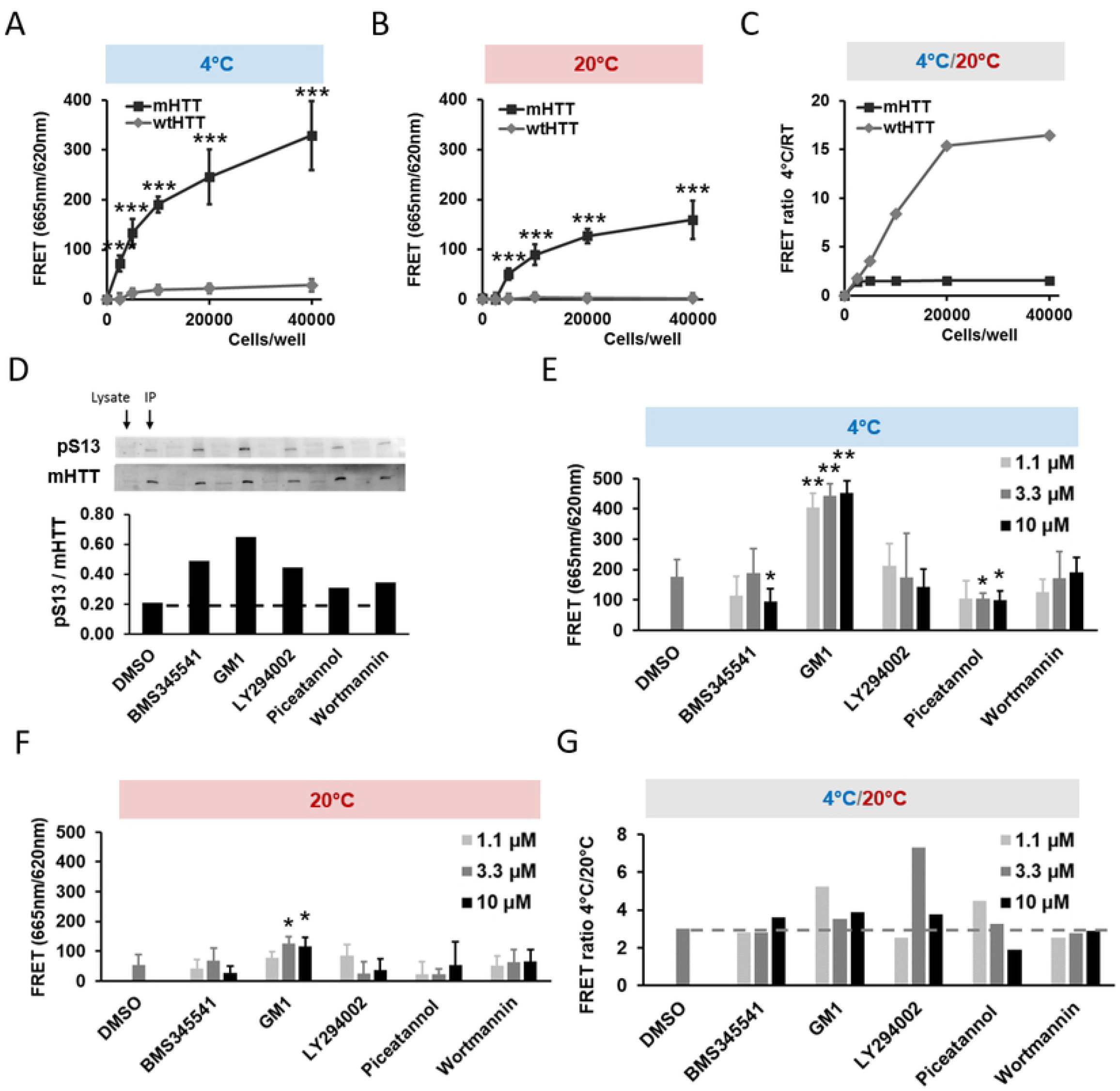
The conformational flexibility of endogenous wtHTT and mtHTT can be distinguished and modified in HD patient-derived human fibroblasts. (A) and (B) Endogenous mHTT and wtHTT expressed by human fibroblasts can be distinguished from each other both at 4°C and 20°C in a cell number-dependent manner. Statistical comparison between cell lines. (C) The 4°C/20°C TR-FRET ratio is higher for the wHTT than mHTT due to the stronger wtHTT signal decrease at 20°C. (D) Immunoprecipitation of HTT followed by Western blotting of HTT and HTT pS13 in HD patient-derived fibroblasts recapitulates that several published small molecules can increase S13 phosphorylation levels. Cells were treated with 10 μM for 16 hours. (E) and (F) mHTT TR-FRET in fibroblasts treated with the compounds in (D) at 4°C and 20°C. Cells were treated with the indicated concentrations for 16 hours. Statistical comparison with DMSO control. (G) Several small molecules increase the HTT 4°C/20°C TR-FRET ratio in HD patient-derived fibroblasts. The dashed line represents the signal strength of the DMSO treatment condition. All data are represented as mean ± S.D. of at least three independent experiments. Statistical significance was calculated using the paired two-tailed Student’s t-test (*=p<0.05; **=p<0.01; ***=p<0.005).

Previous work from several laboratories identified, among other residues, the N-terminal HTT serine 13 (S13) to be sufficient for increasing mHTT flexibility when phosphorylated or replaced by a phosphomimetic aspartic or glutamic acid residue (22,29). Additionally, at least three studies have shown that small molecule compounds, which can positively influence the phosphorylation state of HTT S13 (21–23)_were able to modify mHTT conformation and toxicity in *in vitro* and *in vivo* models of HD. These studies therefore suggest that pharmacological modulation of this residue could potentially modify mHTT flexibility.

We tested if the phosphorylation of endogenous HTT on S13 by previously identified chemical compounds (21,22) could be reproduced and if this PTM could restore the conformational flexibility of mHTT in fibroblast cells. After immunoprecipitation (IP) of HTT with the monoclonal MAB2166 antibody, we were able to detect a mild increase in S13 phosphorylation with the small molecule compounds described in the literature (Fig 2 D). We then tested the effect of these compounds on mHTT flexibility. For LY29002, piceatannol, and ganglioside GM1 we observed an increase in HTT flexibility as demonstrated by the elevated 4°C/20°C TR-FRET ratio (Fig 2 E-G). Despite this, we were not able to establish a clear dose-response relationship for these compounds (Fig 2 E). Together these findings confirm that mHTT PTMs can be modified pharmacologically and that such modifications have the potential to increase the conformational flexibility of mHTT, although effects were modest.

### TR-FRET-based screening can identify modifiers of mHTT flexibility

Having determined that the HTT TR-FRET immunoassay is sensitive enough to identify compounds that can modify mHTT conformation in cells endogenously expressing mHTT, we then used this temperature-sensitive assay in a semiautomated high-throughput screen to identify novel small molecules with the potential to increase mHTT flexibility. The *Z’* values for the miniaturized cell-based assay were 0.36 and 0.53 when wtHTT and mHTT-expressing cell lines were compared at 20°C and 4°C, respectively, indicating consistency and reproducibility across the assays at both tested temperatures (30). We further validated that the half-life of mHTT was sufficiently long to treat cells with compounds for 24 hours without observing decreasing mHTT levels due to transcription or translation inhibition (S2 Fig). The screen was performed in duplicate in 384-well plate format using 3,378 small molecular compounds present in the Sanofi mode-of-action library (Fig 3 A). Compounds were added at a single concentration of 3 μM and incubated with the cells for 24 hours. Next, cells were lysed and the 2B7-Tb/MW1-D2 antibody mix was added. After an overnight incubation, plates were read at 20°C followed by an incubation at 4°C for two hours and a second read-out. The assay was highly reproducible and robust between replicates: At 4°C (Fig 3 B) and 20°C (Fig 3 C) DMSO treated control wells (Fig 3, blue dots) clustered together indicating only a small technical variance (R^2^_DMSO_ = 0.042 and 0.105 at 4°C and 20°C, respectively). In contrast, tested compounds clustered around the diagonal, showing that the compound effects on the TR-FRET signal were reproducible for most of the tested molecules in the two replicates (R^2^_Compounds_ = 0.956 and 0.936 at 4°C and 20°C, respectively). A small number of compounds had strong fluorescent properties and perturbed the FRET signal and were consequently excluded from the analysis. In addition, some compounds reduced the TR-FRET signal equally at 4°C and 20°C which indicated that HTT levels were reduced, potentially through cellular toxicity (Fig 3 B & C). True screen hits (Fig 3, black dots) were defined as compounds that increased the 4°C/20°C TR-FRET ratio in both replicates more than three times the standard deviation of the DMSO control around its median (grey shaded zone) (Fig 3 D). Most compounds considered as hits increased the 4°C/20°C TR-FRET ratio approximately two-fold. Interestingly, these hits had no effect on mHTT flexibility at 4°C and clustered together with the DMSO control (Fig 3 B), while at 20°C most hits were among the lowest scoring compounds (Fig 3 C). A reduction of the signal at 20°C, but not at 4°C excludes a compound-induced effect on HTT concentration, since a reduction of mHTT would lead to lower TR-FRET signals at both temperatures (Fig 1 D & 2 C). Together this indicates that mHTT flexibility can be altered at higher temperatures while at lower temperatures mHTT is too rigid to be positively influenced by the same small molecular compound.

**Fig 3:**
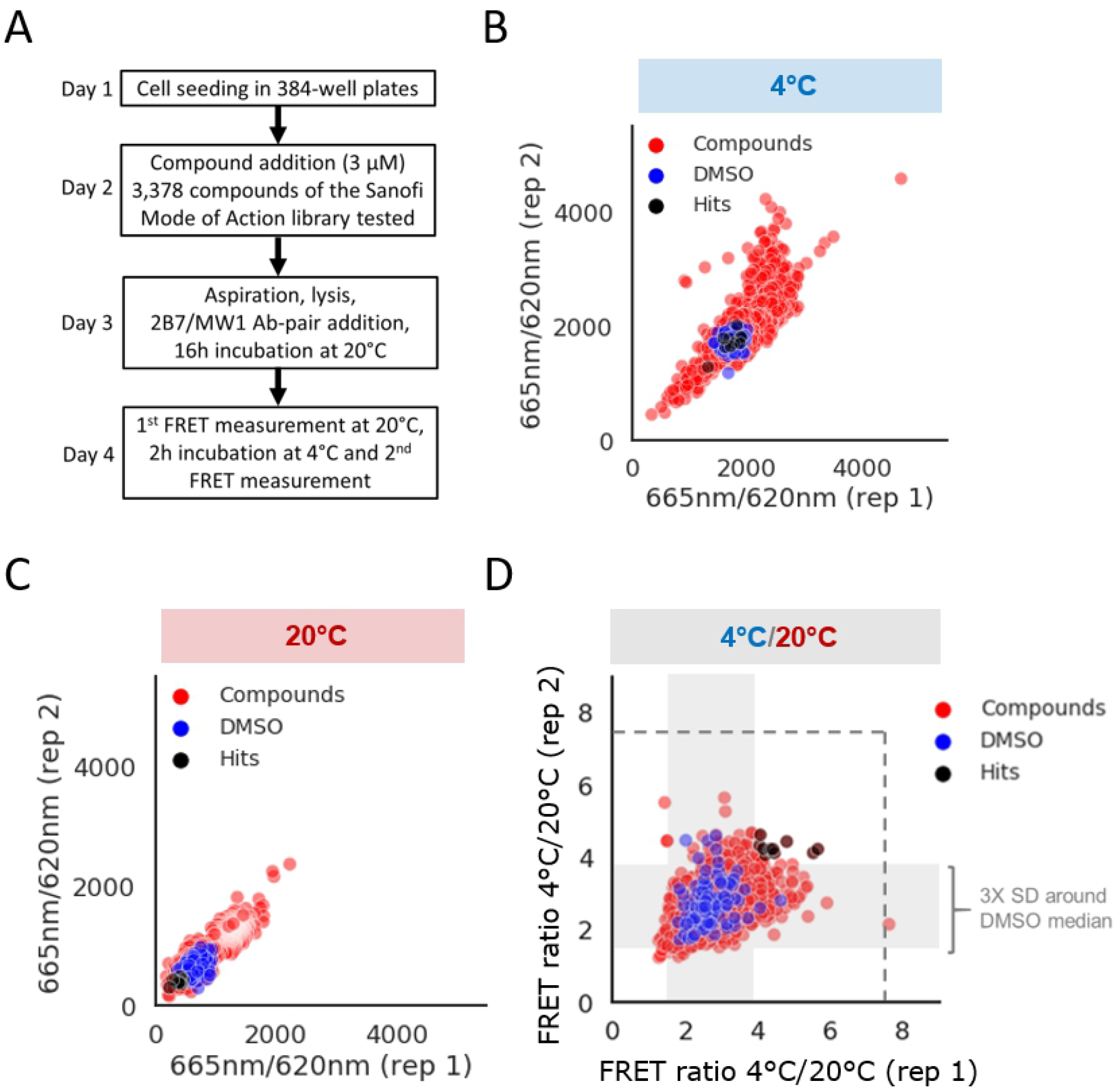
mHTT TR-FRET high-content screening at 4°C and 20°C is technically robust in HD patient-derived fibroblasts. (A) Schematic description of the TR-FRET screening workflow including the compound treatment of living cells. (B) mHTT TR-FRET at 4°C is reproducible and identifies compounds with a broad range of FRET signal modifying activity. Hits described in panel (D) are marked in black. (C) mHTT TR-FRET at 20°C is reproducible and identifies compounds with a broad range of FRET signal modifying activity. Hits described in panel (D) are marked in black and their FRET signal is reduced compared to the DMSO control. (D) Calculating the 4°C/20°C FRET ratio for all tested compounds and DMSO allows to determine the 3X STDEV window (gray area) around the DMSO control median and to identify screening hits. The 4°C/20°C ratio for hits is mainly increased due to their specific reduction of 20°C FRET signal (see panel (C)). The dashed gray line indicates the 4°C/20°C ratio for wtHTT at equal cell concentration (based on data in Figure 2C). R^2^_Compounds_ = 0.956 (4°C) and 0.936 (20°C) indicates biological reproducibility, while R^2^_DMSO_ = 0.042 (4°C) and 0.105 (20°C) indicates technical robustness.

### Identified hits represent multiple modes of action

We selected the ten strongest hits and characterized them further. For reasons of simplicity the compounds are referred to as C1-C10. The majority of the ten compounds have a previously determined biological mode-of-action and defined target protein (Table 1). Compound targets were previously identified by Sanofi in a variety of cell types using a range of biochemical or cellular assays. As a result, some of the identified compounds have multiple targets and additional targets are likely to exist.

**Table 1:**
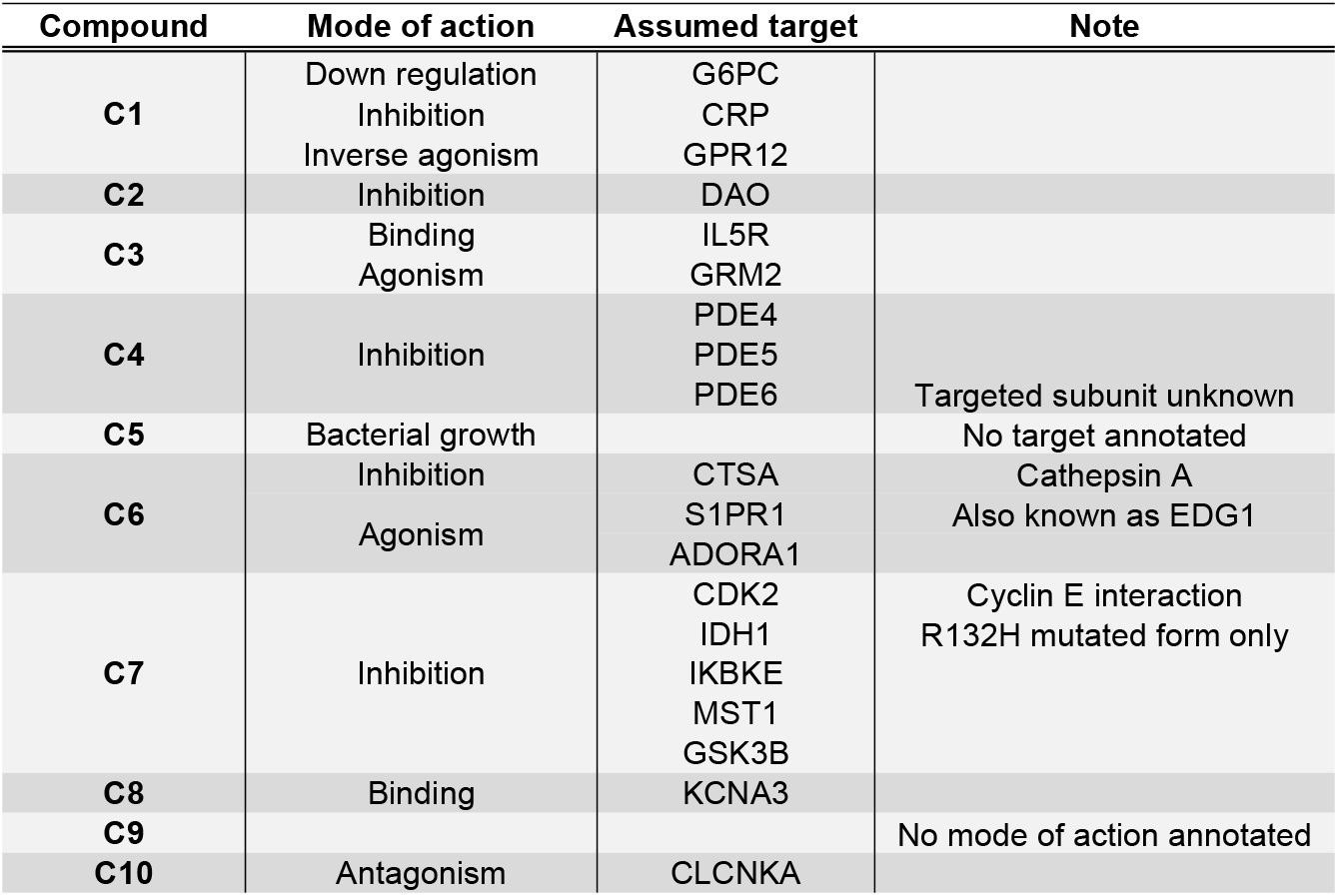
Hit compounds resulting from mHTT TR-FRET high-content screening.

Next, we confirmed the hit compounds by assessing their effect on cell viability, establishing a dose-response relationship, and testing their effect on the phosphorylation of mHTT Ser13 (Fig 4). An ATP concentration-based cell viability assay showed that none of the identified compounds led to a high degree of cell death in the tested concentration range of 0.014 to 30 μM when incubated for 16 hours (Fig 4 A), further confirming that altered TR-FRET signals were not due decreased mHTT caused by cell death. Then we performed HTT TR-FRET with all compounds in the concentration range of 0.014 to 30 μM to identify compounds that had a causal effect on mHTT flexibility. After 16 hours of incubation, we observed that most selected compounds increased the 4°C/20°C TR-FRET ratio 20-30% compared to the DMSO control (Fig 4 B) which was in the same range as observed during initial screening (Fig 3 D), indicating a mild increase in mHTT flexibility. Especially compounds C1, C2, C4, C8, and C9 showed a dose-response relationship towards an increased 4°C/20°C TR-FRET ratio. Effects were generally the highest in the medium tested concentration range between 0.12 and 3.33 μM, which overlapped with the 3 μM used during screening (Fig 3 D & 4 B). Since HTT N-terminal PTMs such as S13 phosphorylation have been described to influence mHTT’s conformational flexibility, we tested whether the identified compounds modified this residue’s phosphorylation relative to the level of HTT. Compound C1 and C10 led to a relative increase of S13 phosphorylation (Fig 4 C). Although, changes in S13 phosphorylation are generally small, this experiment shows that a dual temperature TR-FRET screen can identify small molecules that can potentially modify important PTMs on mHTT and that can in turn modify mHTT conformational flexibility.

**Fig 4:**
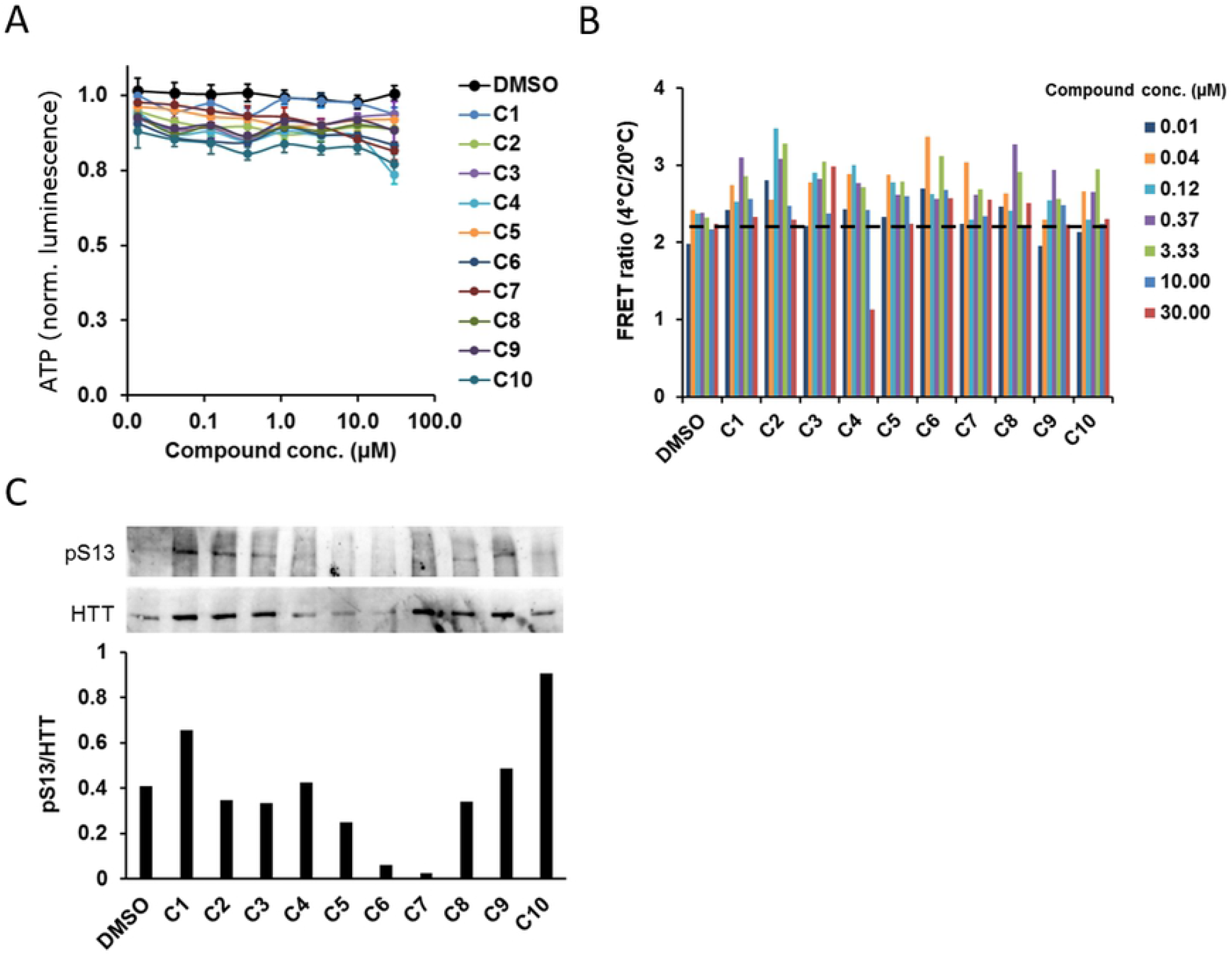
Identified hits have a broad range of activity, consistently increase the 4°C/20°C FRET ratio and partially modify HTT pS13. (A) A cellular ATP assay demonstrates that the 10 strongest identified hits possess a low cellular toxicity. (B) Dose-response testing confirms that compounds increase the 4°C/20°C FRET ratio, but fail to show a dose-dependent effect. For easier comparison with the different treatment conditions, the dashed line represents the average signal strength of all DMSO treatment conditions. DMSO was added at equal volume as DMSO-dissolved compounds. (C) Several compounds alter HTT expression levels, but not specifically pS13 levels.

## Discussion

Glutamine-expanded HTT is regarded as the molecular cause for HD pathophysiology. Reducing mHTT levels in the brain is one strategy to alleviate disease symptoms and severity. Currently, the antisense oligonucleotide (ASO) Tominersen is explored in a clinical trial (ClinicalTrials.gov identifier NCT03761849) and has been shown to reduce mHTT and wtHTT levels in HD patients. However, ASOs have to be repeatedly injected intrathecally, a potentially risky and unpleasant procedure for patients. Furthermore, the depletion of wtHTT might have unintended consequences. Recently, allele selective mHTT depletion has been achieved by a small molecule which both binds to autophagosome protein LC3 and mHTT and induces autophagic clearance (4). Despite such promising results, the functional restauration of mHTT by small molecules rather than its depletion remains a promising and potentially safer alternative. PTMs of the HTT N-terminus regulate HTT structure, aggregation, interactome, toxicity, and cellular properties and have been extensively characterized *in vitro* and are often altered in HD (31–33). Upstream of this molecular cascade lies the polyQ repeat induced alteration of the HTT N-terminal helical conformation. Past work has indicated that mHTT is significantly less flexible which might severely impact its molecular functions (13–16,34). The identification of new drug-like small molecules that can restore mHTT molecular properties, such as helical flexibility, therefore offers and interesting treatment opportunity. However, reproducible high-throughput techniques to identify such candidate molecules are currently not available.

This work demonstrates that high-content screening for modifiers of endogenous mHTT flexibility in HD patient-derived cell lines is feasible with an antibody-based TR-FRET assay. We demonstrate that such an assay is technically sensitive and specific enough in HD patient whole-cell lysates to detect HTT conformational flexibility changes and can be used reproducibly in screening-mode with compound libraries (Fig 5). We detected several small molecular compounds which significantly increase mHTT conformational flexibility, although their exact mechanism of action remains unidentified.

**Fig 5:**
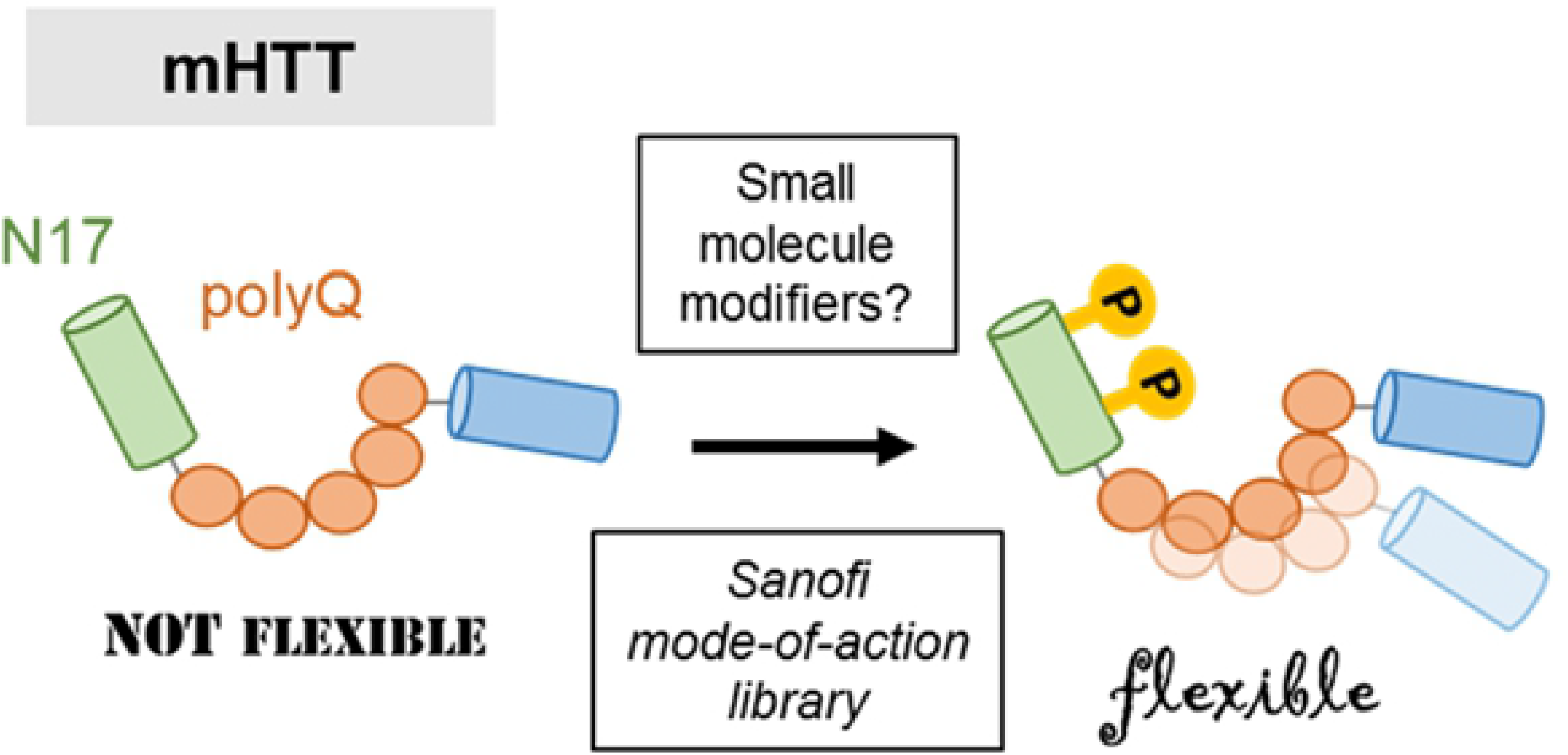
Schematic overview of screening approach. TR-FRET-based screening can identify small molecular compounds that alter mHTT PTMs in living patient-derived human fibroblasts causing a flexibility increase and potentially a functional rescue.

Except for an pS13 HTT antibody, no well-validated antibodies were available to us which can reliably detect other phosphorylated residues in the N17 region which have been previously linked to HTT flexibility, such as threonine 3 (T3) or serine 16 (S16) (17,19,35). In addition, studies have described other N17 PTMs such as acetylation (36,37) or PTMs that are distant from the N17 region and regulate neuronal toxicity (38). It is important to note that the effect of most described PTMs on mHTT conformational flexibility is unknown and needs to be investigated further. The screening approach presented here utilizes endogenous full-length mHTT and therefore has the potential to identify other PTMs that influence the conformational flexibility between the N17 and the polyQ region. Due to the lack of suitable antibodies, in the future detailed mass-spectrometric analysis of HTT fragments will be required to obtain more information about mHTT regulative sites.

As we identified a small number of compounds that seem to influence mHTT flexibility, but also the level of the protein (Fig 3 D & 4 C), our study raises the possibility that dual-action compounds might exist that can be used to lower mHTT levels and in addition partially rescue flexibility. In line with this hypothesis, phosphorylation of certain HTT residues has been found to influence HTT stability. For example, the phosphorylation of mHTT at S434 or S536 decreases proteolysis by caspase 3 and calpain, respectively (39,40). Conceptually similar, phosphorylation at S13 and S16 enhances degradation of both wtHTT and mHTT (29). Identifying new small molecular modulators of mHTT PTMs using conformational flexibility as a read-out might therefore enable the discovery of novel and potentially multifaceted drug targets to treat HD.

## Material & Methods

### Cell lines and cell culture

All used cell lines were obtained from the Venezuelan Huntington Disease Heritable Diseases subcollection of the Coriell Institute, Camden, USA biobank (Table 2). Unless indicated otherwise, cell lines were cultured under standard conditions (37°C and 5% CO_2_) using cell culture medium containing Dulbecco’s Modified Eagle’s Medium (DMEM, Gibco, #105566-016) supplemented with 1X Penicillin/Streptomycin (100X, Gibco, #15140-122) and 15% fetal bovine serum (FBS, Gibco, #16000-044).

**Table 2:** Overview of cell lines used in study.

| Cell line name | Sex | Age at sampling | HD status (polyQ length) |
| --- | --- | --- | --- |
| GM04729 | Female | 39 | Unaffected (Q17) |
| GM09197 | Male | 6 | Affected (Q151) |
| GM04723 | Female | 19 | Affected (Q67) |
| GM04476 | Male | 57 | Affected (Q45) |
| GM04691 | Male | 31 | Affected (Q54) |
| GM05539 | Male | 10 | Affected (Q86) |

### Cell line immortalization

The healthy donor GM04729 and HD patient derived GM04691 fibroblast cell lines were used for screening and immortalized. The primary cell lines were co-transduced with lentiviral vectors expressing hTERT and E6E7 at MOI 40 with 4 μg/ml polybrene.

The expression of immortalization genes was confirmed by RT-PCR. After the immortalization procedure the absence of virus was verified using the Lenti-X p24 Rapid Titer Kit (Takara Bio USA, #632200).

### DNA transfection

DNA plasmids containing HTT with 23 or 73 CAGs were obtained from Coriell Institute, Camden, USA (#CH00022 and #CH00023, respectively). 16 hours prior to transfection wtHTT expressing GM04729 cells were plated at 300×10^3^ cells/well in a 6-well plate. 6 μl Fugene 6 (Promega, #E2691) and 90 μl DMEM medium (without FBS) were mixed and incubated for 5 minutes at RT. 2 μg of the respective DNA plasmid were added to the mix and incubated for another 20 minutes. Next, 100 μl of the transfection mix were added dropwise per well and cells were incubated at 37°C and 5% CO_2_. 16 hours posttransfection, all transfected wells were washed with PBS. Cells were detached using 100 μl Trypsin (Gibco, #25200-056) and 1.9 ml DMEM and collected in Falcon tubes. Cells were then mixed in a two-fold dilution series with GM04729 wtHTT expressing cells in order to achieve the desired DNA plasmid dilution steps. Cells were then incubated another 32h to achieve a total DNA plasmid expression time of 48 h and HTT TR-FRET was performed.

### Huntingtin TR-FRET and data analysis

On Day 1, unless stated otherwise, 50 μl cell culture medium containing a total of 10×10^3^ cells were plated in 384-well plates (Greiner, #781080). Plates were incubated for 24 hours at 37°C and 5% CO_2_. On Day 2, if required, cells were treated with compounds or an equal volume of DMSO (Sigma, #D5879-500ML) for 24 hours. On Day 3, 40 μl of medium was aspirated (rest: 10 μl) and cells were lysed in 10 μl of TR-FRET lysis buffer (10mM Tris pH 7.4 (Sigma, #T1503 and #T3253), 150mM NaCl (Sigma, #S9888-500G), 2% protease/phosphatase inhibitor cocktail (Thermo Scientific, #1861281), 0.6% Igepal (Sigma, #18896-50mL)) and incubated for 1 hour at 20°C under light shaking. 3X antibody stock solutions were prepared containing 0.075 ng/μl 2B7-Tb and 0.75 ng/μl MW1-D2 in TR-FRET antibody buffer (10mM Tris pH 7.4, 150 mM NaCl, 0.25% Igepal, 0.1% Bovine Serum Albumine (BSA, Sigma, #A7030-500G)). 10 μl of the prepared 3X 2B7-Tb/MW1-D2 antibody mix were added to the 20 μl remaining in each well yielding a final antibody concentration of 0.025 ng/μl (total well: 0.75 ng) 2B7-Tb and 0.25 ng/μl (total well: 7.5 ng) MW1-D2. Plates were sealed with Parafilm (Sigma, #P7793-1EA) and incubated at 20°C for 16 hours. Next, plates were read on a PHERAstar plate reader (BMG Labtech) in dual-emission time-resolved fluorescence mode (200 flashes, 60–460 μs integration window, focal height: 8.9 mm, TR-FRET optical module: 337 nm excitation and 665 nm and 620 nm emission passing). After reading, plates were incubated at 4°C for 2 hours and plate reading was repeated using identical settings. For data analysis, first the averages for all technical duplicates were calculated. Second, the 665 nm / 620 nm emission ratios were calculated. Division by the donor emission (620 nm) normalizes for fluorescence artifacts. As a last step, the ratio of the Ab pair only control was subtracted from all protein-containing conditions to correct for background FRET caused by effects unrelated to the target protein (see equation).

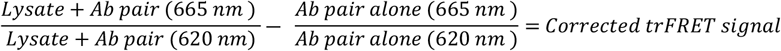

### Immunoprecipitation and Western blotting

In a 10 cm dish cells were incubated with 50 μM GM1 or 10 μM of all other compounds or 0.1% DMSO for 16 hours and then lysed in 1.4 ml Cell Extraction buffer (Invitrogen, #10009222) supplemented with protease/phosphatase inhibitor cocktail (Thermo Scientific, #1861281) and scraped. Lysates were sonicated three times for 30 s each, followed by incubation on ice for 30 minutes. Next, lysates were centrifuged at 10,000 × g for 10 minutes and the supernatant was transferred to a new tube. For the immunoprecipitation of HTT, 400 μl cell lysate were incubated with 4 μl HTT MAB2166 rabbit primary antibody (Merck, #MAB2166) for 2 hours at 4° C on a rotating device. 10 μl of Protein A/G agarose beads (Santa Cruz Biotechnology, #sc-2003) were added to the mix and incubated at 4° C on a rotating device overnight. The immunoprecipitate was collected by centrifugation at 1,000 x g for 30 seconds at 4° C. The supernatant was carefully aspirated and discardedand pellet washed with ice-cold PBS, followed by centrifugation. The pellet PBS wash was repeated two more time. After the final wash, the supernatant was aspirated, discarded and the pellet resuspend in 2X 40 μl of Laemmli electrophoresis sample buffer (Biorad, #161-0737) supplemented with 5% v/v β-mercaptoethanol (Sigma, #M-3148). In a water bath samples were boiled for 2 minutes and 12 ul of each lysate were loaded on two separate gels (Biorad, #4568096), one each required for parallel HTT and pS13 HTT detection, and gels were ran for 60 minutes at constant 200V. Proteins were transferred to a PVDF membrane (Biorad, #1704156) using the high molecular weight option on a Biorad Turbo transfer machine (Biorad) using the high molecular weight option. Blocking was performed in TBS-T + BSA (150mM NaCl (Sigma, #S9625-1KG), 20mM TBS (Thermo Scientific, #28358), 0.1% Tween 20 (Sigma, #P-7949), 5% BSA (Sigma, #A7030-500G)) for 1 hour at 20°C using light agitation. Next, membranes were incubated with HTT (Merck, # MAB2166, Lot: 3286108) or pS13 HTT (Coriell Institute (origin: CHDI), #CH01115, Lot: PGK271101P) antibodies at a dilution of 1/1,000 v/v in blocking buffer at 4°C overnight using light agitation. Membrane washing was performed for 3x 5 minutes in TBS-T using light agitation. Secondary HRP rabbit (Jackson, #711-036-152, Lot: 128838) and mouse (Jackson, #315-036-003, Lot: 127437) antibodies were added at a dilution of 1/1,000 v/v in separate blocking buffers per membrane and incubated at 20°C for 1 hour using light agitation. Membranes were then washed 3x 5min in TBS-T using light agitation. Chemiluminescent substrate (Thermo Fisher Scientific, #3476) was mixed at a ratio of 1/1 according to the manufacturer’s protocol, added to the membranes, and incubated for 5 minutes protected from light at 20°C using light agitation. For imaging, membranes were placed within a transparent plastic on the imaging tray of the Biorad imager and images were taken at sufficient exposure without saturating the image (here: 300 seconds, without pixel binning). Images were exported in TIFF format and pS13 HTT/total HTT ratios were determined using the FIJI/ImageJ Gel Analyzer plugin.

### High-content screening

High-content screening was performed in duplicate and identical to the HTT TR-FRET experiments described above in 384-well format, except for the following changes: Using robotic pipetting, on Day 2 cells were treated with 3 μl/well compounds (3 μM of each library compound in DMSO) or 0.1% DMSO for 24 hours. The two first and last plates were treated with DMSO only to ensure technical robustness and to monitor possible signal fluctuations during the screening procedure. On Day 3 robotic pipetting was used to aspirate the medium and to add lysis buffer and the 2B7-Tb/MW1-D2 antibody mix as described above. Plate sealing was performed manually. On Day 4 plates were placed at 4°C, transported on ice to a PHERAstar plate reader (BMG Labtech) and read-out manually. Data analysis and plate mapping were performed with the Genedata Analyst (Genedata) and TIBCO Spotfire (Perkin Elmer) software packages as well as custom-made KNIME and Python scripts.

### Cell viability assay

On Day 1 GM04691 fibroblasts were counted and diluted to 200×10^3^ cells/ml. 50 μL of each dilution were added per well in 384-well plates yielding 10×10^3^ cells/well. Plates were incubated at 37°C and 5% CO_2_ for 24 hours. On Day 2 5 μl of a 10X concentrated threefold dilution series for each compound were added to the cells, yielding a final 1X concentration of 30 μM to 0.01 μM. Plates were then incubated for another 24 hours. On Day 3 plates were removed from the incubator, equilibrated at 20°C for 30 minutes and 30 μl CellTiter-Glo Luminescent Cell Viability Assay reagent (Promega, #G7573) was distributed into each well. Plates were centrifuged briefly and lightly agitated for 2 minutes at RT. The luminescence signal was allowed to stabilize for 15 minutes and plates were read on a PHERAstar plate reader (BMG Labtech) using the luminescence detection mode. ATP standards were used to estimate the linear measurement range of the assay.

## Acknowledgements

We would like to thank Lee Varban (Sanofi Cambridge, MA, USA) for technical advice concerning HTT TR-FRET in fibroblasts, Robert Godemann (Evotec, Hamburg, Germany) and Doug MacDonald (CHDI foundation) for conceptual project discussions and technical advice, Nicolas Muzet and Rama Heng (Sanofi Strasbourg R&D Center, France) for help in processing compound annotations, and the Coriell Institute biobank for providing the pS13 HTT antibody and HTT DNA plasmids.

## Supporting information

**S1 Fig: Identification of suitable cell lines for screening.** (A) Different primary wt (Q17) and HD patient fibroblast lines were tested in the TR-FRET assay to define which lines would give the best signal to background (S/B) ratio for the the screening. A blank (no cells) control was also used. (B) Western blotting confirmed the expected HTT expression in fibroblast lysates from healthy (wtHTT, 17Q) and heterozygous HD patients (mHTT, 54Q). The difference between wtHTT and mHTT is 37Qs equaling 4.77 kDa and can be detected with a Tris-Acetate gel run for 6 hours at 110V and the ab109115 antibody. For HD patient-derived lysates total protein concentrations ranging from 30 to 5 μg were loaded to highlight size differences of mHTT and wtHTT bands.

**S2 Fig: Determination of HTT protein stability after Cycloheximide (CHX)-induced translation inhibition.** (A) Detection of wtHTT and mHTT using a Tris-Acetate gel and the ab109115 antibody after treatment of GM04691 fibroblasts with the translation inhibitor CHX for the indicated duration. (B) Quantification of wtHTT and mHTT bands indicated in (A). Both wtHTT and mHTT were stable for at least 24h. (C) BCA assay-determined total protein concentration decreases over time. (D) Levels of both wtHTT and mHTT rose relative to total protein.

**S3 File:** Data accompanying Figure 1–4.

